# Suspended and sinking particle-associated microbiomes exhibit distinct lifestyles in the Elbe estuary

**DOI:** 10.1101/2024.10.09.617383

**Authors:** Sven P. Tobias-Hünefeldt, Jason N. Woodhouse, Hans-Joachim Ruscheweyh, Shinichi Sunagawa, Vanessa Russnak, Wolfgang R. Streit, Hans-Peter Grossart

## Abstract

Estuaries are important components of the global carbon cycle; exchanging carbon between aquatic, atmospheric, and terrestrial environments, representing important loci for blue carbon storage and greenhouse gas emissions. Estuarine particles are especially important due to their microbial transformation and vertical/horizontal transport. We used metagenomes and metatranscriptomes to assess changes in microbial community composition and functions across the Elbe estuary over one year, linking changes to dissolved and particulate organic matter. We will be the first to link microbial activity derived from molecular data to particulate and dissolved organic carbon characteristics. There were no microbial species responses to the measured physicochemical and dissolved/particulate organic matter parameters, however Weighted Correlation Network Analyses showed significant spatial microbiome differences linked to the estuarine salinity profile. Meanwhile, suspended and sinking particle fractions did not show any community wide differences, but individual gene analyses revealed clear microbial lifestyle differences. Sinking particle-associated transcripts highly expressed competition and stress-responses genes, while suspended particle-associated transcripts favoured energy acquisition and growth. Transcription patterns further indicate that suspended particles may represent a mitigating influence on methane release via methanotrophy, while both suspended and sinking particles produce methane through the same gene complex. This notion suggests that increased sinking particle abundance, such as under high turbidity conditions, leads to increased methane production. Our findings further imply that urban activities such as dredging may have a high impact on greenhouse gas emissions, and higher suspended-sinking particle ratios may alleviate aquatic-atmospheric methane exchanges. Future studies should explore in detail the underlying mechanisms and controlling variables, in particular when taking predicted global changes into account.

## Introduction

Aquatic environments act as CO_2_ storage and processing centres, production, and utilisation, and transportation hubs (Gao et al., 2022), linking terrestrial and oceanic ecosystems (Regnier et al., 2022), and terrestrial-atmosphere transfers (Liu et al., 2022). Estuaries in particular are critical carbon hotspots with their disproportionate carbon cycling influence (Hutchings et al., 2020). During productive seasons (i.e. spring and summer), estuaries may act as carbon sinks. However, the high CO_2_ production in freshwater areas leads to an overall net CO_2_ production annually (Zimmermann-Timm, 2002). As a result, estuaries have large carbon footprints and influence carbon transport (Khatiwala et al., 2013; Rackley, 2010), playing a key role in greenhouse gas emissions and climate change feedback.

Estuaries are characterised by a distinct salinity gradient and dynamic environmental conditions including freshwater flow and linked marine intrusions which affect biodiversity, heterotrophy rates, food-web efficiency, and seawards nutrient/carbon transport (Chilton et al., 2021; Giblin et al., 2010; Gillanders & Kingsford, 2002; Santoro, 2010). Essentially, all biogeochemical dynamics and carbon exchanges, both atmospheric and seawards, are altered and can lead to eutrophication. Salinity creates a barrier that freshwater organisms cannot overcome, and/or, marine organisms (such as phytoplankton) are found deeper in the estuary where they are sheltered, including in side-channels, where they may be productive. Consequently, phytoplankton bloom rate increases, leading to oxygen depletion following bloom-collapse, with hypoxic areas increasing in prevalence and size over time (Sanders et al., 2023). Oxygen depletion leads to many effects by creating hostile conditions, killing or isolating endemic species, and shifting ecosystem conditions, affecting biogeochemical processes such as carbon cycling.

One of the most common forms of carbon is particulate matter. These particles are major hotspots of activity for microbial degradation, and consumption (Simon et al., 2002; Zimmermann-Timm et al., 2002), linking dissolved and particulate carbon states. They transport carbon between pelagic, benthic, and aquatic environments with sequestration in estuarine and marsh sediments and massive resuspension under high discharge conditions. The Elbe estuary (Germany) is an important system for studying the dynamics of particles and dissolved organic matter in estuarine environments. It is one of the largest estuaries in Europe, supporting a wide range of habitats and species, and is of commercial importance. Numerous attempts have been made to understand the source and fate of inorganic/organic carbon in the Elbe estuary (Matoušů et al., 2018; Tobias-Hünefeldt et al., 2024), although focus was usually on freshwater rather than brackish zones (Amann et al., 2012, 2015; Cisternas-Novoa et al., 2015; Jürgens et al., 1997; Zimmermann-Timm, 2002). Previously, differences between suspended and sinking particles have been identified both in the Wadden Sea, and within the Elbe estuary (Lunau et al., 2004; Tobias-Hünefeldt et al., 2024). Elbe particles differed in density and POC, linking to either marine-like or terrestrial, humic-like DOM. Additionally, Elbe particles were highly variable due to different anthropogenic influences, such as dredging, tidal marsh and harbour exchanges, to seasonal phytoplankton and zooplankton dynamics. This is why previous studies identified significant differences between particles of upper and lower parts of the Elbe estuary.

Bacterial influences play a central role in particle degradation and DOM turnover. Microbial particle degradation is a major source of CO2 (Cai, 2011; Simon et al., 2002) and an important driver of biogeochemical cycles (Jiao et al., 2010; Longhurst & Glen Harrison, 1989) due to the high associated metabolic potential and activity of particle-associated microbes compared to free-living organisms (Grossart et al., 2007; Lyons & Dobbs, 2012; Nguyen et al., 2022; Wang et al., 2024). While estuary effects (i.e. the associated nutrient/salinity gradient) on microbial particle degradation and therefore the carbon cycle have been explored (Lin et al., 2024), studies have yet to integrate physicochemical profiles, microbial community compositions, functional potential, metatranscriptomes, DOM profiles, and particle characteristics. Although studies have identified shifts in the particle-associated community composition and colonisation in response to the estuarine gradient, (Jürgens et al., 1997; Tobias-Hünefeldt et al., 2024; Wörner et al., 2002), the microbiome composition represents only one aspect of the variable factors. Therefore, it is important to understand what microorganisms and genes are present, their metabolic capabilities, and establish links between particle organisms and specific functions to gain a comprehensive understanding of the carbon transfer through the system (Y. Liu et al., 2020; Trevathan-Tackett et al., 2019; Urvoy et al., 2022; Zoccarato & Grossart, 2019).

With this study we sought to explore spatiotemporal gene patterns in the Elbe estuary and compare and contrast the microbial community composition and function of sinking and suspended particles with that of the free-living microbial community, including particulate and dissolved carbon dynamics. We anticipated that particles would have a reduced microbial diversity, due to the short natured presence of particles before disaggregation through sheer-stress, as previously hypothesised (Tobias-Hünefeldt et al., 2024). Particle-associated microbiomes would also be more involved in the breakdown of complex polymers than free-living bacteria. Furthermore, we hypothesised that suspended particles are enriched for enzymes involved in the degradation of algal polymers and proteins, whereas sinking particles contain higher abundances of recalcitrant material degradation enzymes. We expand on previous findings by including the influence of microbes on the Elbe’s biogeochemical carbon cycle and flux across the salinity gradient, and assess osmoregulation genes to identify if the salinity gradient is responsible for the decreased bacterial colonisation rates and survivability (Jurdzinski et al., 2023).

## Materials and Methods

### Sampling

Samples were taken from the River Elbe estuary in the main channel, Germany, in May-21, Jul-21, Feb-22, May-22, Jun-22, and Nov-22 at 5 stations (Mülenberger Loch [53.54907, 9.82338], Twielenfleth [53.60921, 9.56536], Schwarztonnensand [53.71442, 9.46976], Brunsbüttel [53.88742, 9.19429], and Medemgrund [53.8363, 8.88777]) (Figure 1). Samples were taken with a horizontal sampler (Lunau et al., 2004) at a depth of 1 m, due to very high Elbe turbidity that rapidly decreases primary production over depth. Suspended and sinking particles were allowed to vertically separate for 30 minutes, and downstream particle analyses were carried out on each separate fraction. Free-living microbes were captured on 0.22 µm Durapore filters, using filtrate from the particle-associated microbes, captured on 5-µm Durapore filters. All samples were collected in duplicate, unless otherwise stated.

**Figure 1.**
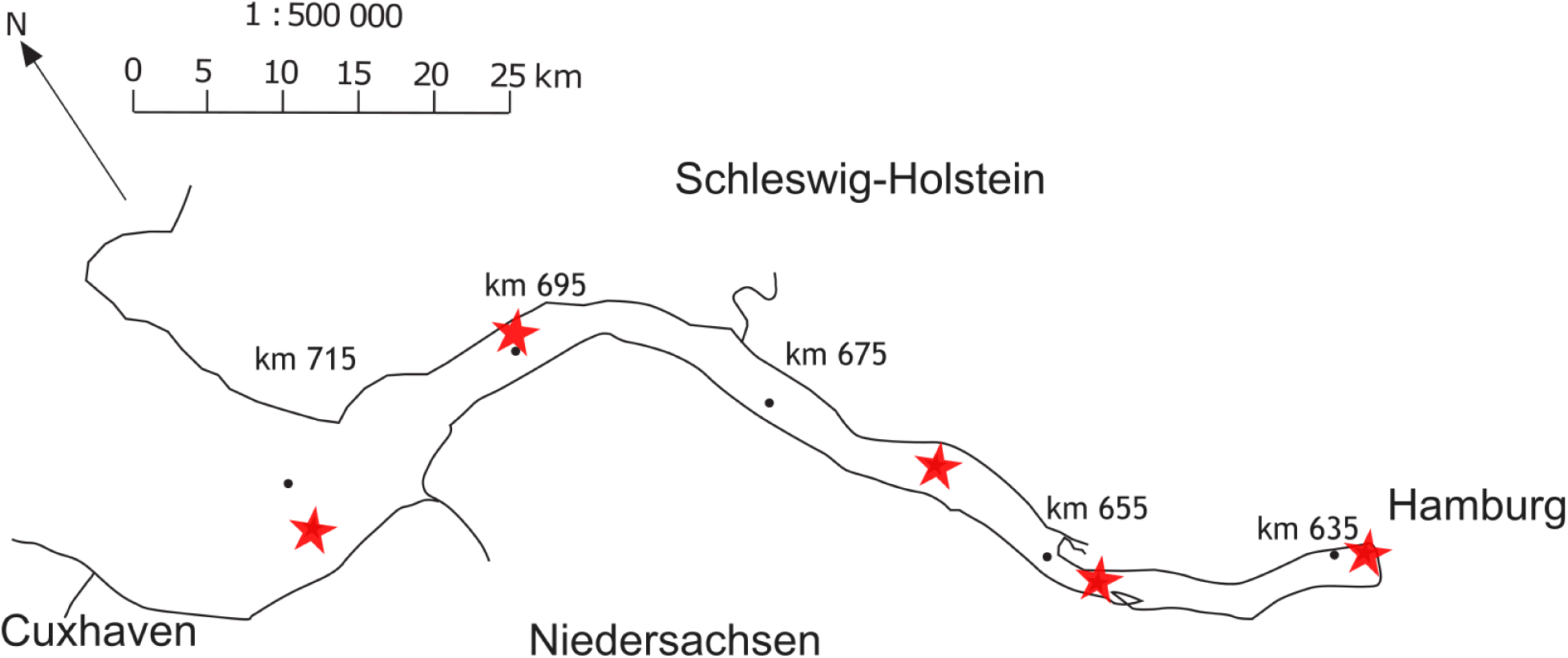
Elbe Estuary map. The map depicts the outline of the Elbe, from Hamburg Harbour to Cuxhaven, with marks of the Elbe length (in kilometre) shown. Sample stations are shown as stars. Particle characteristics were obtained by filtering samples onto pre-weighed and rinsed GF75 filters, followed by another rinsing to remove salts, to determine their particulate matter dry weight and particulate total and organic carbon contents. Before use, filters were pre-combusted for 5 hrs at 400 °C to remove any residual carbon.

### DNA extraction and sequencing

DNA was extracted using the method described in (Nercessian et al., 2005). In brief, cells were lysed with zirconia-silica beads (0.1-1 mm) suspended in cetyltrimethyl ammoniumbromide (CTAB). Sodium dodecyl sulfate and N-Lauroylsarcosine (anion surfactants), and proteinase K and phenol-chloroform-isoamyl alcohol were added. Chloroform-isoamyl alcohol and polyethylene glycol (PEG) was used for DNA purification, and precipitation at 4°C, ethanol washing air-drying and finally dissolving in Tris (Tris-hydroxymethyl-aminomethane). Metagenomic sequencing was performed at Ramaciotti Centre for Genomics (Sydney, Australia) and the Competence Centre for Genomic Analysis Kiel (Kiel, Germany). Samples were prepared for sequencing with the Illumina DNA prep kit, and sequenced on a NovaSeq 6000 platform (Illumina, San Diego, CA, USA). Raw sequences are available on NCBI under BioProject accession number PRJEB54081 (BioSamples: SAMEA110290250 - SAMEA110290357, SAMEA110291994 - SAMEA110292011, and SAMEA112714775 -SAMEA112714862). Sequences were processed as outlined in (Paoli et al., 2022), with metaSPAdes (Nurk et al., 2017), BWA (Li & Durbin, 2009), metaBAT 2 (Kang et al., 2019), Anvi’o (Eren et al., 2015), CheckM (Parks et al., 2015), Prokka (Seemann, 2014), DIAMOND (Buchfink et al., 2021), and KEGG (Kanehisa & Goto, 2000). With an emphasis on processes involved in the processing of monomeric and polymeric organic carbon substrates, we established an extensive genome and gene catalogue to understand shifts in composition and function of free-living, suspended and sinking particle-associated microbiomes.

The CO_2_ and CH_4_ gene database was manually curated based on the Kyoto Encyclopedia of Genes and Genomes (Kanehisa, 2019; Kanehisa et al., 2023; Kanehisa & Goto, 2000), extracting all CO_2_ (C00011) and CH_4_ (C01438) associated genes.

### Physicochemical parameters

Salinity, oxygen, temperature and turbidity were measured with the onboard FerryBox (Petersen et al., 2011). Bulk water was utilised to measure dissolved CO_2_ and CH_4_ concentrations using the headspace technique. In brief, 100 mL of water was collected in a 500 mL syringe, 400 mL of pure N_2_ gas was added and the syringe vigorously shaken for 1 minute. 350 mL of gas were transferred to 1 L Gas Bags and the CO_2_ and CH_4_ concentration measured within 12 hours using an Ultraportable Los Gatos (Los Gatos Research, USA), as described in Kang et al. (2024).

Particulate Organic and Total Carbon (POC/PTC), dissolved organic carbon (DOC), dissolved ammonium, dissolved nitrate, dissolved nitrite, soluble reactive phosphate (SRP), and total dissolved nitrogen, phosphate, and silicate were determined as outlined in Tobias-Hünefeldt et al. (2024).

### Statistical analysis

All figures were created using ggplot2 (version 3.5.1) (Wickham, 2016) and finalised with Inkscape (version 1.3.2) unless otherwise stated. Group differences were assessed with the ANOSIM and PERMANOVA tests with the vegan package (version 2.6-6.1), and ANOVA tests from the stats package (version 4.1.2); eta_squared() from the rstatix package (version 0.7.0) assessed the correlation strength of ANOVA tests. Differences between two group means were assessed with wilcoxon.test() from the stats package, pairwise Wilcoxon tests from the stats vegan package identified pairwise differences from groups. Spearman tests with cor.test() from the stats package assessed correlations between two trends.

Network analyses were carried out with the WGCNA package (version 1.72-5) in R, genes that were excluded from the network are removed from subsequent statistical analyses. Gephi (version 0.10.1) was used to visualise the network. Mantel tests from the vegan package (version 2.6-6.1) were used to compare distance matrices (generated using dist()) between mOTUs, metagenomes, and transcripts per genome, and their network module eigenvalues. Metatranscriptome derived mOTUs could not be utilised, due to the low number of matches between metatranscriptomes and marker genes, therefore all mOTUs are derived from metagenomic data.

All code is available at https://github.com/SvenTobias-Hunefeldt/ElbeMicrobiome.

### Results & Discussion

The Elbe’s prokaryotic microbiome was analysed for its community composition (mOTUs), functional potential (metagenomics), and transcripts per gene (metatranscriptomics). We correlated microbiome characteristics to previously established physicochemical gradients in the Elbe estuary (Tobias-Hünefeldt et al., 2024) for free-living and particle-associated organisms. We identified significant spatial and temporal effects for all microbiome aspects, with temporal differences driving community composition and functional potential while transcripts were primarily driven by spatial factors. Free-living prokaryote community composition and functional potential were driven by temperature, and nitrate determined transcription patterns due to high *de novo* biosynthesis and alternative energy pathways. This also explains the previously identified lack of aromaticity and humic patterns.

We split particle microbiomes into suspended and sinking particles associated. Unlike our initial hypothesis and previous studies in the Wadden Sea (Lunau et al., 2004), we could not detect significant community level differences between sinking and suspended particles. Higher turbulence in the Estuary compared to the Wadden Sea keeping particles in suspension and in constant aggregation-disaggregation. However, individual genes still showed significant differences between the two fractions, with transcription differences indicative of different lifestyles.

### Particle-water column characteristics are spatiotemporally entwined

Overall, we identified significant associations between physicochemical parameters, across particle and water characteristics. These correlations were often intertwined with spatiotemporal characteristics, with future climate dependent effects, such as droughts, increasing CH_4_ release, and the hypoxic area size and possibly duration.

A complete correlation matrix of measured environmental parameters identified several strong correlations (Figure 2), the strongest between POC-PTC and NO_2_ ^-^–NH_4_ ^+^. This study focuses on carbon exchange and particle characteristics, as such increased focus was given to dCO_2_, dCH_4_, TEP, and CSP interactions.

**Figure 2.**
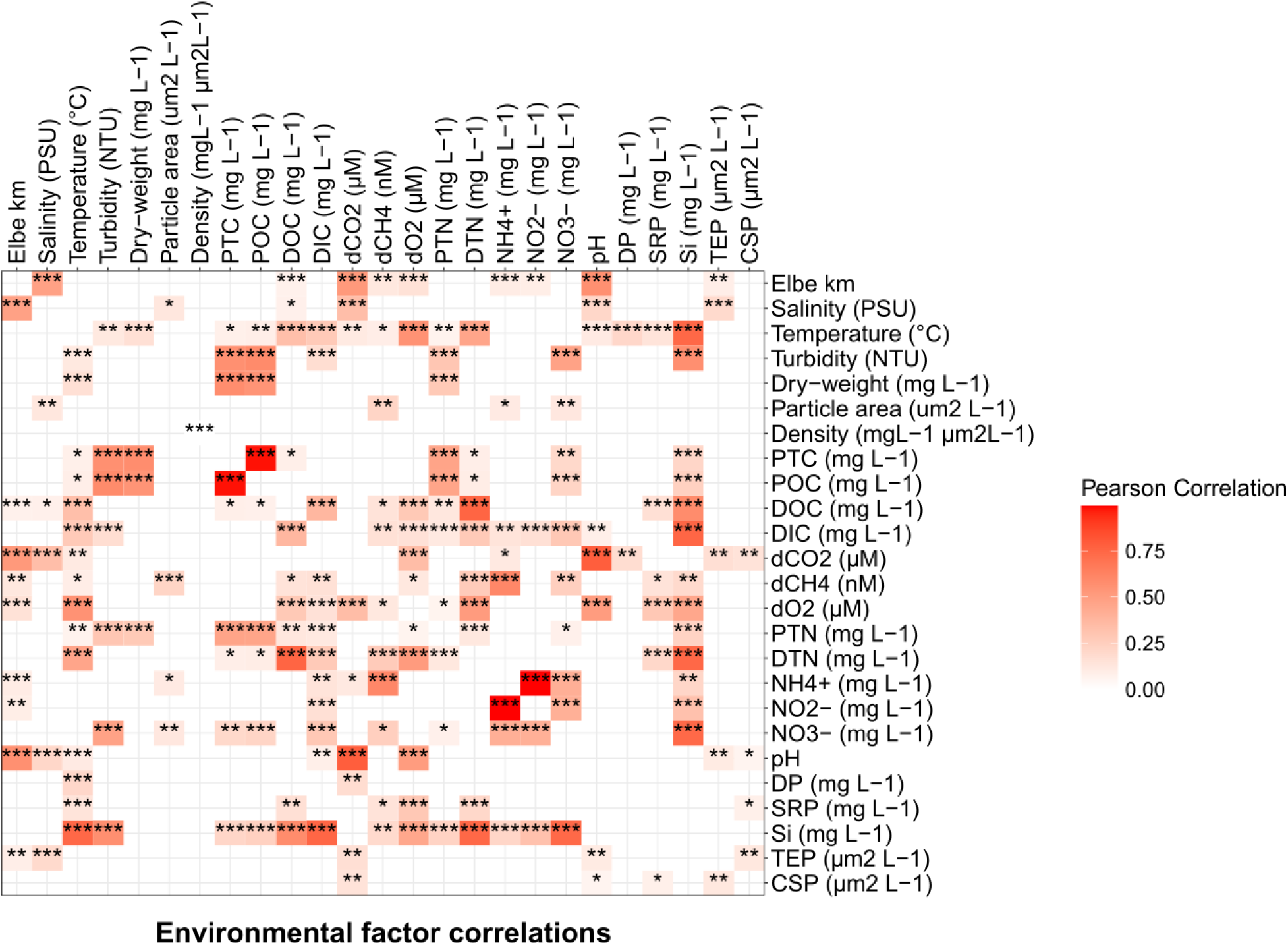
Environmental factor correlations. Colours denote Pearson correlation coefficient between environmental factors, with stars denoting p-value (" " non-significant, * < 0.05, ** < 0.01, **\*** < 0.001).

TEP and CSP were significantly correlated (p < 0.01) to each other, also sharing correlations with pH and CO_2_. However, only TEP correlated with salinity and spatial location, while CSP matched soluble reactive phosphate (SRP) patterns, which may relate to phytoplankton abundance dynamics. TEP formation is pH sensitive (van Pinxteren et al., 2022), and abundance decreases in response to decreases in pH as their structure is disrupted. If only pH was responsible for their abundance, abundance should have increased seawards together with pH (Figure S1), therefore we must conclude that pH is not the sole influence on TEP abundance. It is likely that CSP exhibits a similar effect, with previously identified changes in the stained portion based on acidic vs. neutral pH (Chial & Splittgerber, 1993).

dCO_2_ concentrations have been shown to be dependent on water flow rates in estuaries, and local production and distribution (Raymond et al., 2000). dCO_2_ is a useful parameter to identify activity in an ecosystem, specifically an autotrophic vs. heterotrophic balance. Especially so when paired with dissolved O_2_ concentrations and key carbon fixation gene transcription abundances (Figure S2). Dissolved O_2_ showed significant seawards increases (Spearman, rho = 0.39, p < 0.01), the reverse of dCO_2_ patterns (Spearman, rho = -0.87, p < 0.01), indicative of algal primary production (Figure S3). Seawards stations did not exceed atmospheric CO_2_ concentrations, meanwhile Mühlenberger Loch did. Suggestive of more net heterotrophic than autotrophic activities releasing CO_2_ and absorbing dO_2_; whereas, at seawards stations atmospheric CO_2_ was absorbed due to autotrophic activities. Indicating high autotrophic vs. heterotrophic activities in freshwater vs. brackish Estuary regions. This has been studied extensively, especially with the increased duration and depth of the hypoxic area near Hamburg Harbour (Sanders et al., 2023). Key carbon fixation pathways complemented these findings. Key carbon fixation transcript abundances increased seawards at all time points, with November 22 showing a large increase after Hamburg Harbour in carbon fixation genes, matching dCO_2_ concentrations (Figure S2). The harbour represents the highest CO_2_ and CH_4_ values and the hypoxic area would expand further downstream under decreased flow rate conditions as those predicted by increased droughts under climate change conditions. Although dredging in November may have resulted in increased dCH_4_ levels in the Harbour region. Meanwhile, increased floods increase sediment resuspension and heterotrophic activity in the Elbe (Huang et al., 2015; Kerner, 2000a). In the context of TEP and CSP concentrations we can conclude that higher abundances in autotrophic environments are due to phytoplankton production. Other salinity related changes have previously been discussed (Tobias-Hünefeldt et al., 2024). CSP is primarily composed of protein compounds, such as amino acids. Many of these amino acids are synthesised via phosphorylated pathways, therefore elevated SRP concentrations lead to an increase of amino acid production, with increased primary production also driving increased protein synthesis.

Most physicochemical parameters significantly correlated with temperature, a good proxy for season in the Elbe (ANOVA, eta^2^ = 0.90, p < 0.01). Temperature was also commonly correlated with physicochemical parameters, due to large seasonal differences between them. Nitrification processes affect pH in the Elbe, with large temporal and spatial influences (Amann et al., 2012), with nitrification decreasing pH, due to the reduction of free protons when producing nitrate and nitrite. Nitrification processes which have been shown as seasonally influenced via remineralisation/nitrification activity near Hamburg Harbour, and upstream phytoplankton growth (Sanders et al., 2018).

Dissolved CH_4_ revealed significant seasonal differences. One of the potential reasons for the high February-22 concentrations can be attributed to meltwater resuspending sediments and degassing sediment CH_4_. Biomarkers for methane production vs. consumption are mcrA and pmoA (Friedrich, 2005; McDonald & Murrell, 1997), reflecting methane flux within the environment (Kong et al., 2019; Nazaries et al., 2013). We could not identify native water column production, as classical methanogenesis by methanogenic Archaea typically occurs in soils and sediments rather than oxic water columns (Blake et al., 2020; Jabłoński et al., 2015; Sherry et al., 2016). Methanotrophy on the other hand could be identified in the transcripts, specifically from Gammaproteobacteria (Ga0077536, Methylococcaceae, and Methylomonadaceae). Transcriptional patterns showed highest activity in free-living organisms, with high transcription rates seawards in June 22, and in brackish areas in November 22. Meanwhile, particle-associated pmoA transcripts decreased seawards, identical to all metagenome fractions (Figure 3). Water retention increases free-living microbe abundance (Luef et al., 2007), and therefore droughts in the Elbe would decrease dCH_4_ uptake across the estuary, releasing more CH_4_ into the atmosphere.

**Figure 3.**
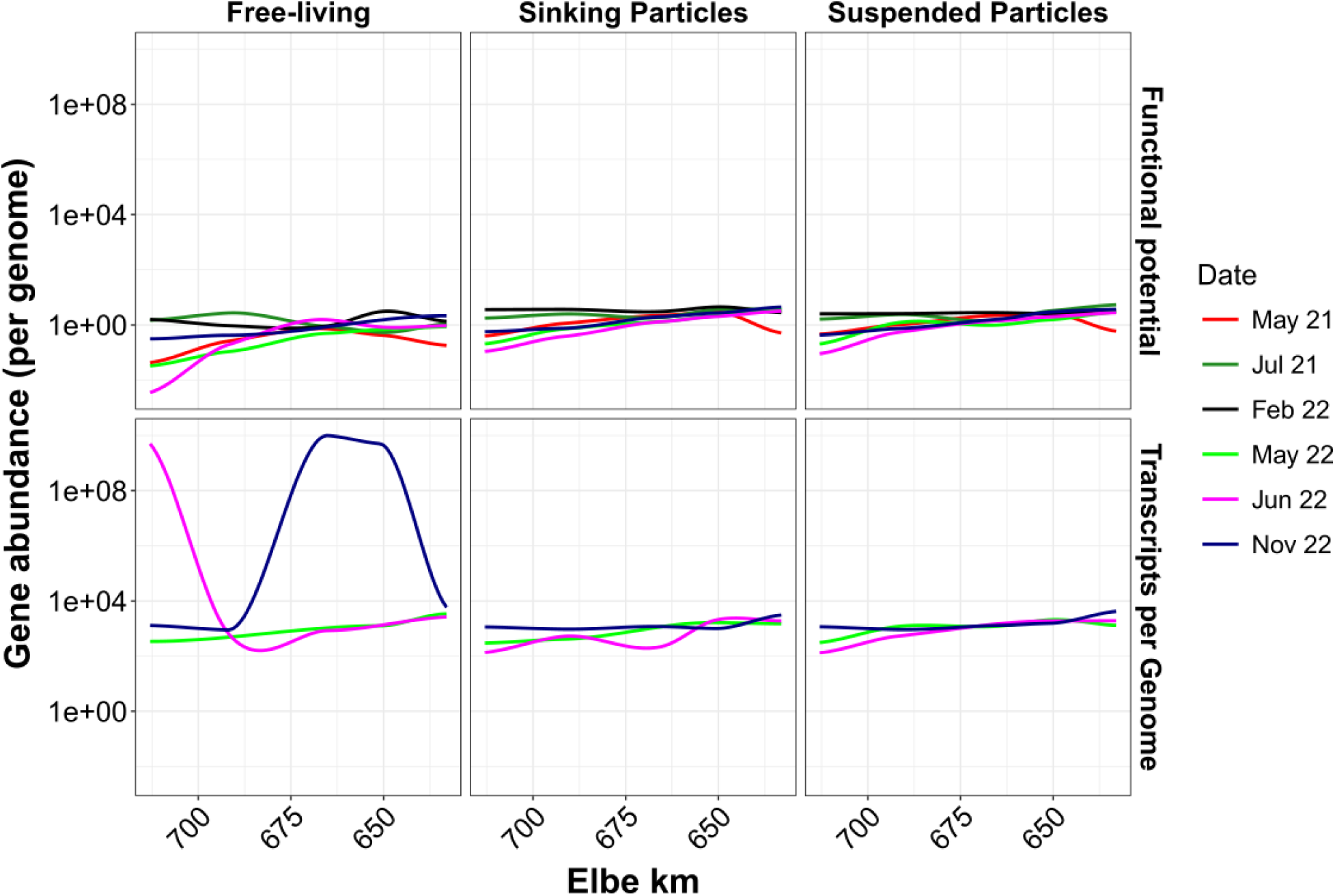
pmoA abundance in the Elbe estuary. The mean of two samples pmoA functional potential and transcripts per genome shown across the Elbe Estuary with colours depicting sample dates. pmoA abundance has been taxonomically corrected against amoA. Fractionated particle-associated and free-living abundances are shown separately.

Overall, we identified significant associations between physicochemical parameters, across particle and water characteristics. These correlations were often intertwined with spatiotemporal characteristics, with future climate dependent effects, such as droughts, increasing CH_4_ release, and the hypoxic area size and possibly duration. This has various knock-on effects such as increased fish-stress (Koll et al., 2024), and the carbon transport efficacy (Li et al., 2024).

### Lifestyle defines particle-associated functional potential and activity rather than community level differences

While community aspects between suspended and sinking particles could be ascertained, individual genes did show significant differences between the two particle fractions. We initially hypothesised that sinking materials would have increased abundances of aromatic compound degradation genes due to a high abundance of resuspended sediments, while the suspended particle microbiome would express genes targeting more labile material due to high abundances of fresh organic matter, such as proteins. As the Elbe is highly turbulent, we expected stochastic particle settlement, with high particle turnover. Although community composition and functional potential may not significantly differ, the transcription patterns would represent particle fraction dependent lifestyles, with suspended particles containing labile material degradation genes.

As we expected, functional potential showed minimal lifestyle differences (Figure 4). Sinking particles contained a significantly increased abundance of *rbcL* (chloroplast gene, involved in photosynthesis and usually in chloroplast DNA (Klein et al., 1994; Nurhasanah et al., 2019))*, pheC* (phenylalanine biosynthesis and upregulated under cold tolerant conditions (Cheng et al., 2023; Zhao et al., 1992))*, csiD* (only expressed under carbon starvation conditions (Marschall et al., 1998))*, oadA* (upregulated in response to citrate and essential for its uptake (Blancato et al., 2008; Repizo et al., 2013)) and *DDC* (converts tryptophan to tryptamine, linking primary and secondary metabolic pathways (Goddijn et al., 1994)). Meanwhile, suspended particles showed higher abundances of *ubiA* (ubiquinone biosynthesis that expresses under stress conditions, specifically low oxygen and carbon (Suzuki et al., 1994))*, alsD* (internal cell pH buffering (Renna et al., 1993))*, bioU* (biotin production, and essential carboxylation, decarboxylation, and transcarboxylation cofactor (Sirithanakorn & Cronan, 2021))*, menA* (menaquinone biosynthesis, important for anaerobic electron transport systems (Suvarna et al., 1998)), and *GGCX* (the only known vitamin K post-translational modification, produces γ-carboxyglutamate, a vital precursor to many proteins including those involved in cellular growth, survival, and signalling (Shearer & Newman, 2014; Shearer & Okano, 2018)). Both suspended and sinking particles show predispositions to genes that are upregulated under stress, whether carbon starvation or anaerobic conditions. Based on these genes we could not identify significant life-style differences, unlike in the transcription patterns. However, it does show that particle-association in the Elbe is fraught, and stress responses are required for survival. However, suspended and sinking particles are under unique stressors.

**Figure 4.**
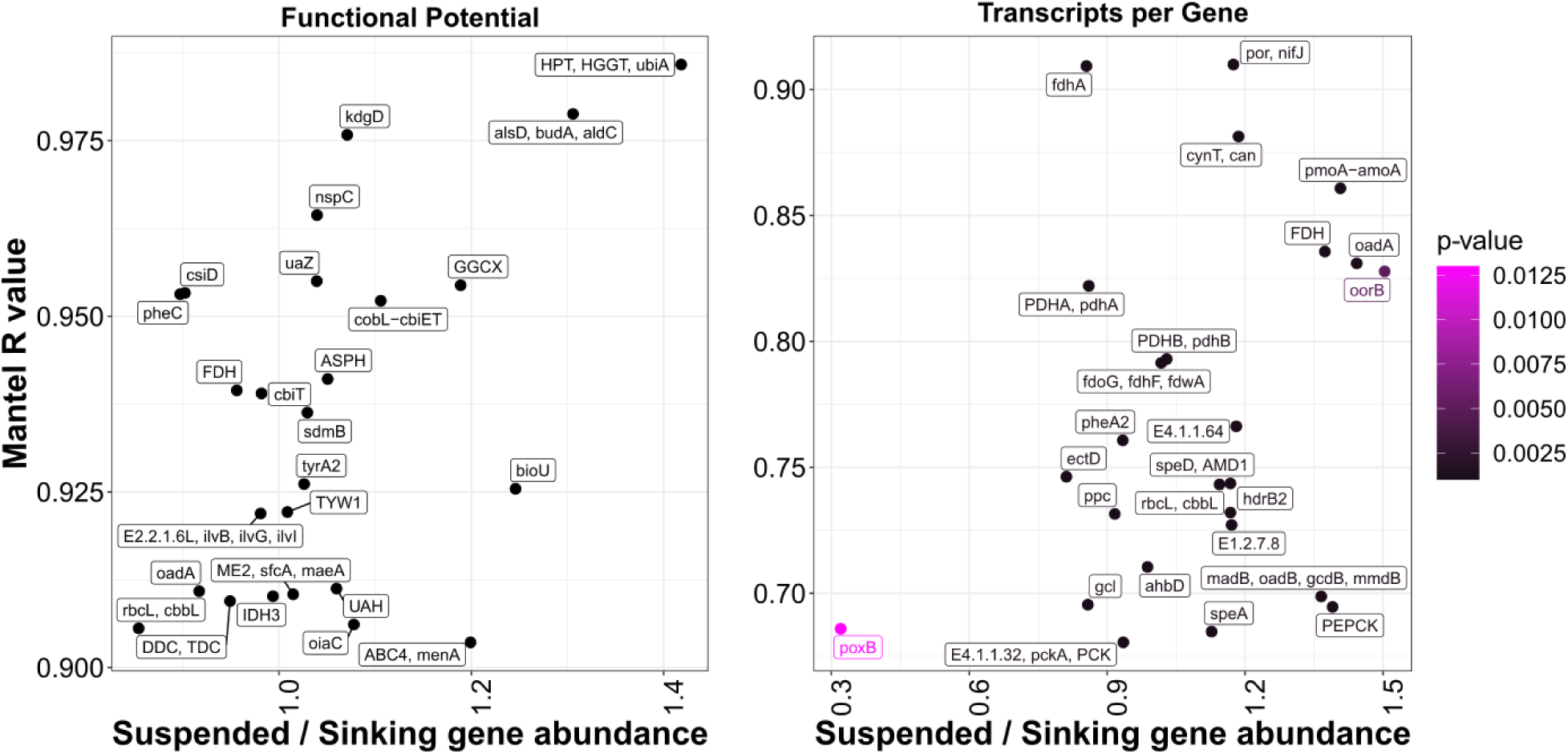
Lifestyle differences between suspended vs. sinking particle-associated functional potential and transcripts per genome. Only the top 25 functional potential genes and transcripts per gene are depicted. Colour denotes p-value, with the y axis representing the Mantel R value, and x-axis the ratio between suspended to sinking abundance.

Individual carbon gene transcript analyses showed that there were significant differences between suspended and sinking particles, indicative of lifestyle. Genes on sinking particles were upregulated in response to stress, including general stress (*poxB* (Weber et al., 2005)), salinity (*ectD* (García-Estepa et al., 2006)), nutrient starvation (*pdhA* (Betts et al., 2002)), and antioxidants (*gcl* (Sigler et al., 1999; Zhang et al., 2020)). Meanwhile, suspended particles show upregulation of PEPCK, downregulated in response to DNA damage and *oorB*, downregulated under hyperosmotic stress. The unequal stress response suggests that sinking particles are under more competition and nutrient limitations, with the expression of *gcl* suggesting predation via phagocytosis (e.g. due to Amoeba (Villalba et al., 2022; Vogel et al., 1980)).

Unlike our predictions, based on anoxia identified in particles, especially under hypoxic conditions (Kerner, 2000b), an aversion to anaerobic conditions on sinking particles was identified, with *pdhA* downregulated under anaerobic conditions, and *poxB* requiring aerobic conditions to produce acetate (Lorquet et al., 2004). While on suspended particles *oadA*, which requires anaerobic conditions, is significantly upregulated. However, when assessing functional potential, *oadA* was significantly increased on sinking particles, showing that while functional potential favours sinking particles, responses to conditions show that *oadA* is upregulated on suspended particles. Showing how both methods can work together.

Sinking microbiomes showed increased stress responses, while suspended microbiomes expressed genes associated with growth and energy acquisition. At the community level, transcription patterns were not closely correlated to POC concentrations unlike community composition and functional potential, due to the organisms preference for dissolved nitrogen sources, such as nitrate, which are present at high concentrations in the Elbe estuary. However, in the event of nitrogen limitations the community is capable of POC degradation.

The top 5 upregulated suspended particle-associated genes are predominantly related to energy acquisition, with fermentation and other carbon acquisition strategies through *oadA*, *poxB*, PEPCK, and *pmoA*. Meanwhile, the top 5 transcript up-regulations associated with sinking particles are represented with only *pdhA*, which is itself upregulated under nutrient starvation conditions. As *pmoA* requires CH_4_, it suggests that suspended particles are involved in alleviating CH_4_ release, whereas sinking particles only produce CH_4_. This should be further explored in future studies.

Overall, while functional potential comparisons did not reveal lifestyle differences, stressors differed between suspended and sinking particles. Transcription patterns, on the other hand, revealed high competition and stress responses in aerobic conditions, while suspended particle communities upregulate energy acquisition and growth pathways under both aerobic and anaerobic conditions.

### Spatiotemporal contributions to microbiome composition and carbon processing

No richness differences between the distinct particle fractions could be identified, only free-living functional potential gene richness was significantly lower than either particle fraction (Wilcoxon, p < 0.01, Figure 5). Meanwhile, transcriptional profiles and community composition profiles showed similar species and active gene numbers across free-living, and suspended and sinking particle-associated microbiomes. These findings support the hypothesised high aggregation and disaggregation rates, with no deterministic microbiome selection due to particle characteristics, or genome streamlining present. Attributed to high turnover rates and the need to continuously recolonise particles. Additionally, findings suggest that gene richness of active genes does not differ between microbiome fractions, with the same number of genes active in each fraction.

**Figure 5.**
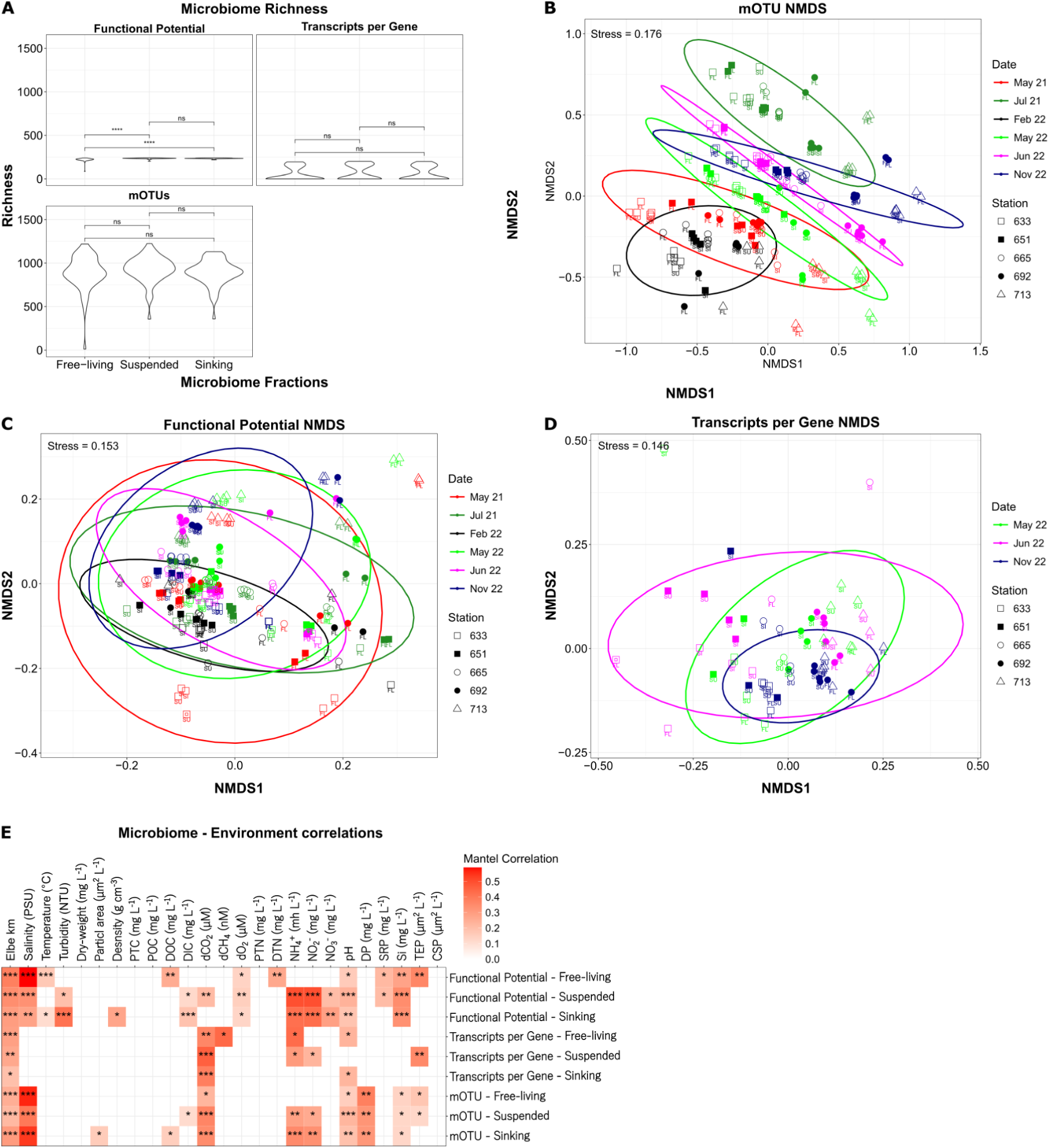
Microbiome aspects in the ecosystem. A violin plot A) showing richness of the same microbiome aspects in the same fractions, pairwise comparisons are made with Wilcoxon tests. NMDs plots of microbiome aspects (B: mOTUs, C: Functional Potential, D: Transcripts per Gene,), colours represent sample dates, and shapes sample stations along the Elbe Estuary. Labels below sample points represent the microbiome fractions, free-living (FL), suspended and sinking particle-associated (SU and SI). Outliers have been removed. E: separated microbiome fraction (free-living, suspended- and sinking-particle associated) composition, functional potential, and transcripts are correlated with a Mantel test (in red). P-value is denoted as stars for A) and E) (" " and "ns" : non-significant, * < 0.05, ** < 0.01, * < 0.001).

NMDS analyses showed significant differences between free-living and particle-associated mOTUs, CO_2_ and CH_4_ related functional potential, and transcription patterns (Figure 5; ANOSIM, R > 0.07, p < 0.01; PERMANOVA, R > 0.06, p < 0.01). During pairwise PERMANOVA analyses no community level differences between the two particle fractions were identified (ANOSIM & PERMANOVA, p > 0.05), while differences between either particle fraction and the free-living microbiome were significant (ANOSIM & PERMANOVA, p < 0.05). Much like particle characteristics and microbiome richness, differences can be attributed to the Elbe’s high shear stress conditions leading to continuous particle turnover, and high mixing between re-suspended sediment and freshly generated material (Tobias-Hünefeldt et al., 2024).

Spatiotemporal differences dominated microbiome dynamics, with sample date dominating microbiome composition and functional potential (ANOSIM, R > 0.37, p < 0.01; PERMANOVA, R^2^ > 0.31, p < 0.01), with all individual timepoints differing significantly (p < 0.05). Meanwhile, transcripts showed weaker date correlation (ANOSIM, R 0.05, p = 0.04; PERMANOVA, R^2^ = 0.04, p = 0.11), where instead sample site played a larger role (ANOSIM, R = 0.13, p < 0.01; PERMANOVA, R^2^ = 0.09, p < 0.01). These site differences were driven by differences between the 2 seawards and three most freshwater sites, also true for microbiome composition and functional potential (p < 0.05).

We identified a significant correlation between the Elbe’s particle characteristics, and community composition (Mantel, R = 0.11, p < 0.01), functional potential (Mantel, R = 0.16, p < 0.01) and transcriptions per gene (Mantel, R = 0.19, p = 0.02). Samples have been retained in their free-living and sinking/suspended particle-associated dynamics (Figure 5). Focusing on carbon relevant physicochemical parameters we found that TEP interacted with the free-living community functional potential and community composition, and suspended particle community composition and transcriptional activity (Figure 5; Mantel, R = 0.17-0.35, p < 0.05). We used Mantel differences to explore whether the environment is “selecting” or the microbiome is generating/degrading the measured physicochemical parameters. The relationships we have identified are all positive, therefore selection and generation are the most likely relationships.

TEP correlations do not represent spatiotemporal differences, such as salinity changes, due to transcriptional profile correlations, which were absent from salinity. Instead, the identified microbiome aspects are likely most involved in TEP generation. Another strong possibility is the utilisation of TEP by these communities. It has been shown that while phytoplankton may generate TEP, other prokaryotes may modify TEP without decreasing its abundance (Stoderegger & Herndl, 1998), resulting in a positive relationship between them, without directly generating TEP. Only free-living functional potential, suspended particle microbiome aspects, and free-living and suspended microbiome aspects significantly correlated to TEP abundance. We suggest that only the free-living aspect correlated due to the creation of EPS, including TEP, in their metabolic potential as suspended particle-associated organisms instead focus on the utilisation of labile material, and production patterns could not be identified due to their lower abundances. Meanwhile, free-living organisms produce an extracellular matrix, including TEP, to generate an advantageous environment (Netrusov et al., 2023). Similarly, a strong correlation with salinity may have been a confounding factor, and further studies must identify the true cause. It is also possible that non EPS producing organisms outnumber EPS producing organisms on particles, and therefore we do not see the same TEP relationship between particle-associated organisms and TEP abundances (Jayathilake et al., 2017). We suggest that the relationship between suspended transcription profiles and TEP abundance is due to the microbiome’s ability to modify TEP, similarly to the functional potential. Meanwhile, free-living organisms are producing EPS, and sinking microbiomes are focused on recalcitrant material degradation, as discussed in the particle lifestyle section. mOTU patterns are a combination of both of the previous points, with only sinking microbiomes not significantly related to TEP abundances. Likely due to its lack of TEP modification, or generation.

On the other hand, we did not identify significant microbiome-CSP abundance correlations (Mantel, p > 0.05). Suggesting that the protein fraction captured in CSP may not play a role in community composition, potential, or activity, but previous evidence argues against this (von Jackowski et al., 2020). Instead, a more likely scenario includes rapid and continuous CSP turnover, as suggested previously (Tobias-Hünefeldt et al., 2024).

Dissolved CO_2_ was the 3^rd^ most universally related physicochemical parameter (after station km and pH), interacting with suspended particle functional potential, and all microbiome compositions and transcription profiles. Meanwhile, dCH_4_ only significantly interacted with free-living transcriptional profiles. dCO_2_ represents one of the core metabolic components, i.e., heterotrophy and primary production, and thus a transcription profile correlation is expected. We have identified three different regions of dCO_2_ concentrations, Mühlenberger Loch-Twielenfleth, Schwarztonnensand-Brunsbüttel, and Medemgrund. Previous studies have identified significant activity changes over the Elbe estuary, matching the same dCO_2_ pattern (van Beusekom et al., 2021). However, the Schwarztonnensand-Brunsbüttel region represents consumption minima, with the Mühlenberger Loch-Twielenfleth representing the maxima. Therefore, the dCO_2_ concentrations represent the heterotrophy-primary production balance in each region, meaning that the Medemgrund region represents the region with the highest primary production, most likely due to a marine influence. This will be further explored with a network analysis.

Dissolved methane (dCH_4_) concentrations were highest in Mühlenberger Loch, a location known for its longer retention of particulate matter, however, we did not correlate dCH_4_ concentrations with any particulate microbiome aspect. Instead, dCH_4_ correlations could only be identified in the free-living transcriptional profile. Methane is predominantly produced in low oxygen environments, and only recently has oxic methane production been discussed. This process may be occurring in the Elbe estuary as dCH_4_ concentrations were at their peak in Mühlenberger Loch when oxygen was similarly at its peak. Further studies are required to definitely identify the source of dCH_4_ in the Elbe, as previous studies identified significantly contributing terrestrial CH_4_ sources, sediments, side-channels, and anoxic Harbour waters (Borges et al., 2018; Y. Li et al., 2021, 2021; Steinsdóttir et al., 2022; Wells et al., 2020). We did not identify significant methanogen patterns in the Elbe estuary that would explain measured dCH_4_ concentrations.

To identify what microbiome aspects were related we ran a series of Weighted Correlation Network Analysis (WGCNA) on all identified mOTUs, MAG associated genes, and transcripts (Figure 6). Contextualising mOTU clustering with the associated MAG if available for its functional potential and transcriptional profile.

**Figure 6.**
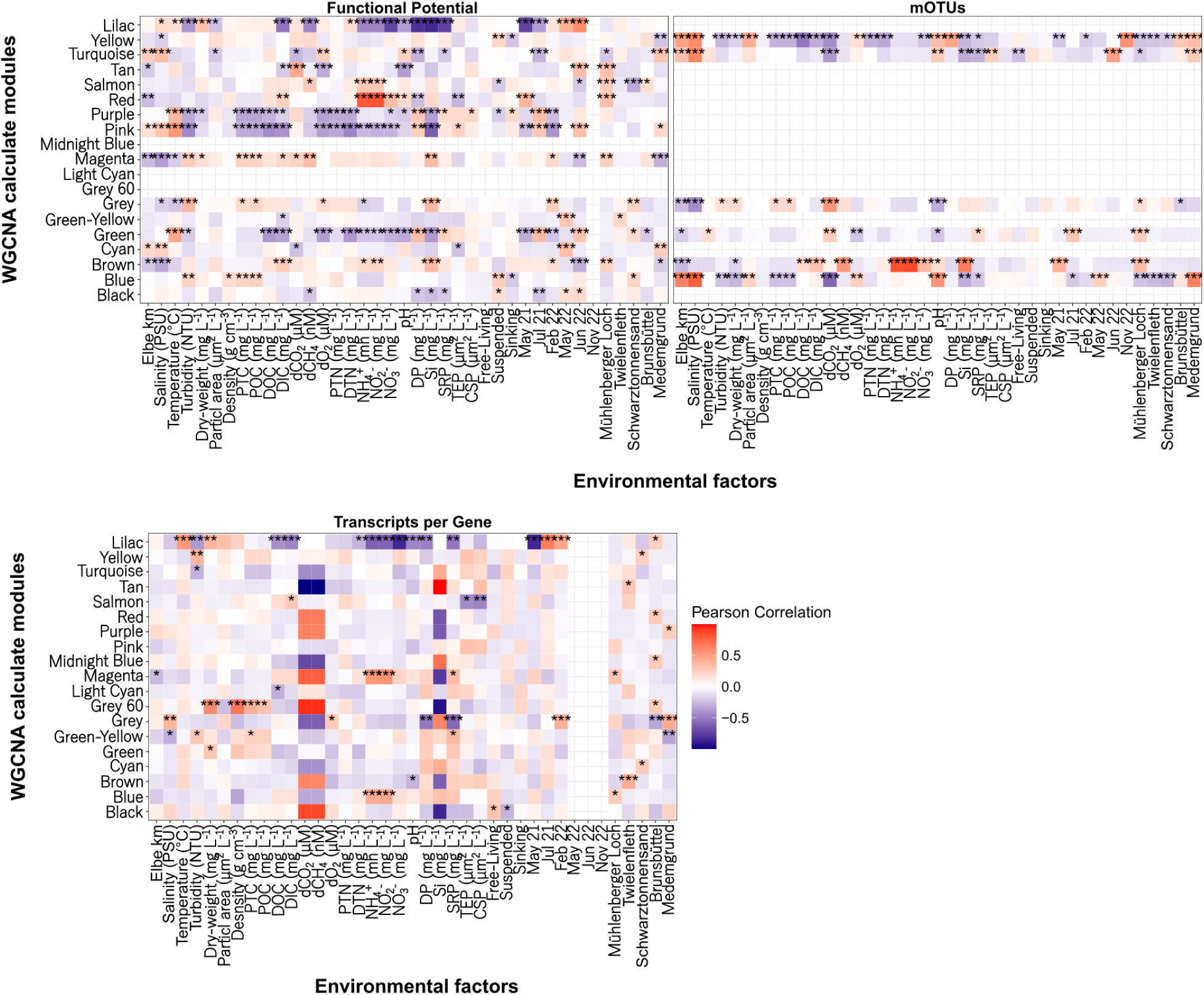
WGCNA network modules generated from microbiome aspects. mOTU, functional potential, and transcript per gene profiles underwent Weighted Correlation Network Analysis (WGCNA). Colours represent the direction and strength of the pearson correlation, while P-value is denoted as stars (" ": non-significant, * < 0.05, ** < 0.01, * < 0.001). WGCNA modules between community aspects are independently calculated.

Carbon processing genes were not specifically grouped into their own modules, with the highest carbon processing metagenome module only containing 5.18% carbon genes (out of 502 module 2 genes). CH_4_ had a maximum of 0.44% in module 2. Transcript network modules were disproportionately influenced by 1-2 samples (Figure S4), making the network unreliable and prone to bias so will not be further discussed. Most carbon processing genes were identified in modules that contained large numbers of genes expressed at low abundances. Such as transcripts per gene module 1 containing 169 CO_2_ genes, out of 4910 total genes (3.44%). Suggesting that the modules containing carbon genes relate to core metabolic functions, rather than specific, or environmental responses, especially as those genes were generally related to core functions such as the TCA cycle. Instead specific responses were identified in other network modules.

Similarly, methanotrophs did not group into their own network, even though we identified modules associated with CH_4_ changes (Figure 6). The module MAGs were instead made up of Bacteroidota, Planctomycetes, Alphaproteobacteria, Pseudomonadota, and Verrucomicrobia. The same occurred with metagenome and transcriptome modules, where we could not identify significant correlations between gene and CH_4_ profiles, specifically we could not identify a significant correlation between *mcrA* and CH_4_. Therefore, we suggest that methane production is limited, or below detectable rates in the Elbe estuaries water column, and instead sediment production dominates.

We identified a salinity dependent mOTU cluster (Figure 6, module 1 / turquoise) where 22.9% of the unique mOTU taxonomies contained osmoregulation associated genes, allowing for increased salinity tolerance (Table S1). The module additionally corresponded to TEP abundance concentrations. It has been shown that increased EPS concentrations alleviate salinity (Mukherjee & Samaddar, 1997), of which TEP is a component. However, TEP has previously been shown to decrease seawards, most likely due to increased sinking rates (Tobias-Hünefeldt et al., 2024), due to salinity promoting TEP aggregation (Meng & Liu, 2016). Therefore, we suggest that taxa in this module are not directly correlated to TEP abundance, but include indirect salinity effects, such as salinity effects on TEP.

A second salinity correlated module was mOTU module 2 (blue), which showed significant increases under the most saline conditions, specifically in Medemgrund in May-22, during the lowest flow rate. 30% of the mOTU taxa contained osmoregulation associated genes, which would allow for saline tolerance. The module is made up of organisms that originate in ocean conditions, such as Opitutales (Choo et al., 2007), Pelagibacterales (Morris et al., 2002), and Flavobacterales (Seo et al., 2017), which have been found in high abundance in coastal sites before. However, the module also includes microorganisms such as Nanopelagicales which are more commonly found in freshwater environments (Neuenschwander et al., 2018), although they have been identified in brackish waters (Hugerth et al., 2015; Mehrshad et al., 2016; Salka et al., 2014).

The two salinity associated modules show that salinity is a significant driving factor, with connections between microbiome aspects and due to physicochemical correlations, may provide false positive associations between microbiome aspects and other physicochemical parameters. Salinity is also not always the deciding factor for taxonomic distributions. mOTU module 3 (brown) showed the highest affiliation with nitrogen compounds, NH_4_, NO_2_, and NO_3_. This module correlated with the Harbour area (Mühlenberger Loch), during low flow rates (May 21) when slow passage through the Harbour area allows for the accumulation of nitrogen compounds in the water column. Identified mOTUs included Bacteroidota, Planctomycetes, Alphaproteobacteria, and Verrucomicrobiae. Only Alphaproteobacteria (*Aquidulcibacter)* contained osmoregulation genes (*envZ*), so it is likely that further increased salinisation due to saline water intrusion to the Elbe estuary would expand their area, affecting nitrogen compound concentrations, specifically through the increased abundance of NO and N_2_O processing genes (Figure S5), although further research would be required.

Overall, we identified significant spatial differences in the microbiome, rooted in functional potential and transcript effects. Most significantly, were the effects of salinity and differences between the Harbour and mostly brackish environment. With nitrogen compounds also playing a large role in all community aspects.

## Conclusion

The Elbe’s microbial community showed significant correlations to physicochemical parameters. However, different microbiome aspects showed varying correlation strengths to measured physicochemical characteristics, across particle and water characteristics. Our findings show future climate dependent effects, such as increased CH_4_ release, and increased hypoxic area size and possibly duration, with important knock-on effects such as decreased fish populations and carbon transport efficacy. The microbial community was also heavily dependent on nitrogen concentrations, for all community aspects, and should therefore be examined in more detail.

While we could not determine fraction-related differences in microbiome composition and functional potential, transcription patterns revealed lifestyle differences. Sinking particle-associated transcripts revealed competition and stress-responses in aerobic conditions, while suspended particle-associated transcripts were associated with energy acquisition and growth in both aerobic and anaerobic conditions. Transcription patterns also suggest that suspended particles may represent a mitigating influence on methane release via its utilisation, while both suspended and sinking particles produce methane through the same gene complex.

Our findings indicate that increased methane production may be a result of increased sinking particle abundance. Especially relevant in high turbidity conditions such as the Elbe estuary where urban activities, i.e. dredging, may impact greenhouse gas emissions, with higher suspended-sinking particle ratios decreasing atmospheric methane emissions from estuaries. Future studies should explore the underlying mechanism and controlling variables in greater detail, especially when considering predicted global changes.

## Acknowledgements

This project was funded by the Deutsche Forschungsgemeinschaft (DFG, German Research Foundation) as part of the project ‘Biota-mediated effects on Carbon cycling in Estuaries’ (407270017/RTG2530) and the Bundesministerium für Bildung und Forschung (BMBF, Federal Ministry of Education and Research) as part of the project ‘Blue-Estuaries’ (03F0864C).

Uta Mallok and Monika Degebrodt measured DOC and POC from provided samples, respectively. We thank Helmholtz-Zentrum Hereon and the Ludwig Prandtl Boat Crew for their help during sampling.

The authors have no relevant financial or non-financial interests to disclose.

Author contributions: Hans-Peter Grossart, and Sven P. Tobias-Hünefeldt came up with the study design. Material preparation, data collection and analysis were performed by Sven P. Tobias-Hünefeldt. Vanessa Russnak performed data collection and analysis. Jason Woodhouse, Hans Joachim Ruscheweyh, and Shinichi Sunagawa performed data analysis. The first draft of the manuscript was written by Sven P. Tobias-Hünefeldt and all authors commented on previous versions of the manuscript. All authors read and approved the final manuscript.

