## Supplementary material for "Suspended and sinking particle-associated microbiomes exhibit distinct lifestyles in the Elbe estuary": Table S1

| DESCRIPTION | KO | Gephi_reassigned | Module Colour ID | Phylum | Class | Order | Family | Genus | Species |
|---|---|---|---|---|---|---|---|---|---|
| opuBD: osmoprotectant transport system permease protein | K05846 | 0 | grey | Acidobacteriota | Blastocatellia | Pyrinomonadales | Pyrinomonadaceae | OLB17 | |
| opuC: osmoprotectant transport system substrate-binding protein | K05845 | 0 | grey | Actinobacteriota | Actinomycetia | Actinomycetales | Microbacteriaceae | Microbacterium | Microbacterium ginsengisoli |
| opuA: osmoprotectant transport system ATP-binding protein | K05847 | 0 | grey | Actinobacteriota | Actinomycetia | Actinomycetales | Microbacteriaceae | Microbacterium | Microbacterium ginsengisoli |
| opuBD: osmoprotectant transport system permease protein | K05846 | 0 | grey | Actinobacteriota | Actinomycetia | Actinomycetales | Microbacteriaceae | Microbacterium | Microbacterium ginsengisoli |
| opuA: osmoprotectant transport system ATP-binding protein | K05847 | 0 | grey | Actinomycetota | Acidimicrobiia | UBA5794 | JAENVV01 | JAENVV01 | JAENVV01 sp905480085 |
| opuC: osmoprotectant transport system substrate-binding protein | K05845 | 0 | grey | Actinomycetota | Acidimicrobiia | UBA5794 | JAENVV01 | JAENVV01 | JAENVV01 sp905480085 |
| opuBD: osmoprotectant transport system permease protein | K05846 | 0 | grey | Actinomycetota | Acidimicrobiia | UBA5794 | JAENVV01 | JAENVV01 | JAENVV01 sp905480085 |
| opuA: osmoprotectant transport system ATP-binding protein | K05847 | 1 | turquoise | Actinomycetota | Actinomycetia | Actinomycetales | Microbacteriaceae | Pontimonas | |
| opuC: osmoprotectant transport system substrate-binding protein | K05845 | 1 | turquoise | Actinomycetota | Actinomycetia | Actinomycetales | Microbacteriaceae | Pontimonas | |
| opuBD: osmoprotectant transport system permease protein | K05846 | 1 | turquoise | Actinomycetota | Actinomycetia | Actinomycetales | Microbacteriaceae | Pontimonas | |
| opuC: osmoprotectant transport system substrate-binding protein | K05845 | 2 | blue | Actinomycetota | Actinomycetia | Actinomycetales | Microbacteriaceae | Pontimonas | Pontimonas sp017852615 |
| opuA: osmoprotectant transport system ATP-binding protein | K05847 | 2 | blue | Actinomycetota | Actinomycetia | Actinomycetales | Microbacteriaceae | Pontimonas | Pontimonas sp017852615 |
| opuBD: osmoprotectant transport system permease protein | K05846 | 2 | blue | Actinomycetota | Actinomycetia | Actinomycetales | Microbacteriaceae | Pontimonas | Pontimonas sp017852615 |
| opuBD: osmoprotectant transport system permease protein | K05846 | 0 | grey | Bacteroidota | Bacteroidia | Cytophagales | Cyclobacteriaceae | ELB16-189 | |
| opuBD: osmoprotectant transport system permease protein | K05846 | 5 | green | Cyanobacteria | Cyanobacteriia | Cyanobacteriales | Microcystaceae | Microcystis | |
| opuBD: osmoprotectant transport system permease protein | K05846 | 5 | green | Cyanobacteriota | Cyanobacteriia | PCC-6307 | Cyanobiaceae | Cyanobium | |
| opuBD: osmoprotectant transport system permease protein | K05846 | 1 | turquoise | Desulfobacterota_B | Binatia | HRBIN30 | JAGDMS01 | | |
| opuA: osmoprotectant transport system ATP-binding protein | K05847 | 1 | turquoise | Desulfobacterota_B | Binatia | HRBIN30 | JAGDMS01 | | |
| opuA: osmoprotectant transport system ATP-binding protein | K05847 | 1 | turquoise | Desulfobacterota_B | Binatia | UBA1149 | CAITLU01 | | |
| opuBD: osmoprotectant transport system permease protein | K05846 | 1 | turquoise | Desulfobacterota_B | Binatia | UBA1149 | CAITLU01 | | |
| opuA: osmoprotectant transport system ATP-binding protein | K05847 | 0 | grey | Planctomycetota | Phycisphaerae | Phycisphaerales | SM1A02 | VGXO01 | |
| opuA: osmoprotectant transport system ATP-binding protein | K05847 | 1 | turquoise | Planctomycetota | Planctomycetia | Pirellulales | Pirellulaceae | JABAAE01 | JABAAE01 sp905480545 |
| opuA: osmoprotectant transport system ATP-binding protein | K05847 | 1 | turquoise | Planctomycetota | Planctomycetia | Planctomycetales | Planctomycetaceae | SXXK01 | |
| envZ: two-component system, OmpR family, osmolarity sensor histidine kinase EnvZ | K07638 | 2 | blue | Proteobacteria | Alphaproteobacteria | Pelagibacterales | Pelagibacteraceae | Pelagibacter | Pelagibacter ubique |
| envZ: two-component system, OmpR family, osmolarity sensor histidine kinase EnvZ | K07638 | 4 | yellow | Proteobacteria | Alphaproteobacteria | Pelagibacterales | Pelagibacteraceae | SYDM01 | SYDM01 sp008638125 |
| envZ: two-component system, OmpR family, osmolarity sensor histidine kinase EnvZ | K07638 | 1 | turquoise | Proteobacteria | Alphaproteobacteria | Rhodobacterales | Rhodobacteraceae | LGRT01 | LGRT01 sp001642945 |
| envZ: two-component system, OmpR family, osmolarity sensor histidine kinase EnvZ | K07638 | 2 | blue | Proteobacteria | Alphaproteobacteria | Rhodobacterales | Rhodobacteraceae | Planktomarina | Planktomarina temperata |
| osmB: osmotically inducible lipoprotein OsmB | K04062 | 0 | grey | Proteobacteria | Gammaproteobacteria | Burkholderiales | Burkholderiaceae | Polynucleobacter | Polynucleobacter difficilis |
| opuBD: osmoprotectant transport system permease protein | K05846 | 0 | grey | Proteobacteria | Gammaproteobacteria | Burkholderiales | Burkholderiaceae | Polynucleobacter | Polynucleobacter difficilis |
| envZ: two-component system, OmpR family, osmolarity sensor histidine kinase EnvZ | K07638 | 0 | grey | Proteobacteria | Gammaproteobacteria | Burkholderiales | Burkholderiaceae | SYFN01 | |
| envZ: two-component system, OmpR family, osmolarity sensor histidine kinase EnvZ | K07638 | 2 | blue | Proteobacteria | Gammaproteobacteria | Burkholderiales | Methylophilaceae | BACL14 | BACL14 sp000168995 |
| opuA: osmoprotectant transport system ATP-binding protein | K05847 | 2 | blue | Proteobacteria | Gammaproteobacteria | Pseudomonadales | Pseudohongiellaceae | UBA9145 | UBA9145 sp001438145 |
| envZ: two-component system, OmpR family, osmolarity sensor histidine kinase EnvZ | K07638 | 3 | brown | Pseudomonadota | Alphaproteobacteria | Caulobacterales | TH1-2 | Aquidulcibacter | |
| opuBD: osmoprotectant transport system permease protein | K05846 | 0 | grey | Pseudomonadota | Alphaproteobacteria | Micropepsales | Micropepsaceae | CAIYRG01 | |
| opuA: osmoprotectant transport system ATP-binding protein | K05847 | 0 | grey | Pseudomonadota | Alphaproteobacteria | Micropepsales | Micropepsaceae | CAIYRG01 | |
| envZ: two-component system, OmpR family, osmolarity sensor histidine kinase EnvZ | K07638 | 4 | yellow | Pseudomonadota | Alphaproteobacteria | Pelagibacterales | Pelagibacteraceae | Pelagibacter | Pelagibacter sp001438335 |
| envZ: two-component system, OmpR family, osmolarity sensor histidine kinase EnvZ | K07638 | 4 | yellow | Pseudomonadota | Alphaproteobacteria | Pelagibacterales | Pelagibacteraceae | Pelagibacter | Pelagibacter sp008638165 |
| envZ: two-component system, OmpR family, osmolarity sensor histidine kinase EnvZ | K07638 | 2 | blue | Pseudomonadota | Alphaproteobacteria | Pelagibacterales | Pelagibacteraceae | Pelagibacter | Pelagibacter sp016778675 |
| envZ: two-component system, OmpR family, osmolarity sensor histidine kinase EnvZ | K07638 | 4 | yellow | Pseudomonadota | Alphaproteobacteria | Pelagibacterales | Pelagibacteraceae | Pelagibacter | Pelagibacter sp017880265 |
| envZ: two-component system, OmpR family, osmolarity sensor histidine kinase EnvZ | K07638 | 2 | blue | Pseudomonadota | Alphaproteobacteria | Pelagibacterales | Pelagibacteraceae | Pelagibacter | Pelagibacter ubique |
| envZ: two-component system, OmpR family, osmolarity sensor histidine kinase EnvZ | K07638 | 4 | yellow | Pseudomonadota | Alphaproteobacteria | Pelagibacterales | Pelagibacteraceae | SYDM01 | SYDM01 sp008638125 |
| envZ: two-component system, OmpR family, osmolarity sensor histidine kinase EnvZ | K07638 | 2 | blue | Pseudomonadota | Alphaproteobacteria | Puniceispirillales | Puniceispirillaceae | UBA3439 | UBA3439 sp016778825 |
| envZ: two-component system, OmpR family, osmolarity sensor histidine kinase EnvZ | K07638 | 0 | grey | Pseudomonadota | Alphaproteobacteria | Rhizobiales | Aestuariivirgaceae | Anderseniella | |
| osmB: osmotically inducible lipoprotein OsmB | K04062 | 1 | turquoise | Pseudomonadota | Alphaproteobacteria | Rhodobacterales | Rhodobacteraceae | Boseongicola | Boseongicola sp905479635 |
| envZ: two-component system, OmpR family, osmolarity sensor histidine kinase EnvZ | K07638 | 1 | turquoise | Pseudomonadota | Alphaproteobacteria | Rhodobacterales | Rhodobacteraceae | Boseongicola | Boseongicola sp905479635 |
| envZ: two-component system, OmpR family, osmolarity sensor histidine kinase EnvZ | K07638 | 1 | turquoise | Pseudomonadota | Alphaproteobacteria | Rhodobacterales | Rhodobacteraceae | RFZI01 | |
| envZ: two-component system, OmpR family, osmolarity sensor histidine kinase EnvZ | K07638 | 2 | blue | Pseudomonadota | Alphaproteobacteria | Rhodobacterales | Rhodobacteraceae | UBA10365 | UBA10365 sp003536295 |
| envZ: two-component system, OmpR family, osmolarity sensor histidine kinase EnvZ | K07638 | 4 | yellow | Pseudomonadota | Alphaproteobacteria | Rhodobacterales | Rhodobacteraceae | UBA5972 | |
| envZ: two-component system, OmpR family, osmolarity sensor histidine kinase EnvZ | K07638 | 4 | yellow | Pseudomonadota | Alphaproteobacteria | Rhodospirillales_A | Casp-alpha2 | UBA4479 | |
| osmB: osmotically inducible lipoprotein OsmB | K04062 | 0 | grey | Pseudomonadota | Gammaproteobacteria | Burkholderiales | Burkholderiaceae | Polynucleobacter | Polynucleobacter sp903944255 |
| opuC: osmoprotectant transport system substrate-binding protein | K05845 | 0 | grey | Pseudomonadota | Gammaproteobacteria | Burkholderiales | Burkholderiaceae_A | UBA2463 | UBA2463 sp945901825 |
| opuA: osmoprotectant transport system ATP-binding protein | K05847 | 0 | grey | Pseudomonadota | Gammaproteobacteria | Burkholderiales | Burkholderiaceae_A | UBA2463 | UBA2463 sp945901825 |

| | | | | | | | | | |
|---|---|---|---|---|---|---|---|---|---|
| opuBD: osmoprotectant transport system permease protein | K05846 | 0 | grey | Pseudomonadota | Gammaproteobacteria | Burkholderiales | Burkholderiaceae_A | UBA2463 | UBA2463 sp945901825 |
| envZ: two-component system, OmpR family, osmolarity sensor histidine kinase EnvZ | K07638 | 5 | green | Pseudomonadota | Gammaproteobacteria | Burkholderiales | Burkholderiaceae_B | CAIKVZ01 | |
| envZ: two-component system, OmpR family, osmolarity sensor histidine kinase EnvZ | K07638 | 0 | grey | Pseudomonadota | Gammaproteobacteria | Burkholderiales | Burkholderiaceae_B | CAISIP01 | |
| osmY: hyperosmotically inducible periplasmic protein | K04065 | 0 | grey | Pseudomonadota | Gammaproteobacteria | Burkholderiales | Burkholderiaceae_B | CAISIP01 | |
| opuA: osmoprotectant transport system ATP-binding protein | K05847 | 2 | blue | Pseudomonadota | Gammaproteobacteria | Burkholderiales | Burkholderiaceae_B | Limnohabitans_A | LimnohabitanA sp021300495 |
| opuBD: osmoprotectant transport system permease protein | K05846 | 2 | blue | Pseudomonadota | Gammaproteobacteria | Burkholderiales | Burkholderiaceae_B | Limnohabitans_A | LimnohabitanA sp021300495 |
| opuC: osmoprotectant transport system substrate-binding protein | K05845 | 2 | blue | Pseudomonadota | Gammaproteobacteria | Burkholderiales | Burkholderiaceae_B | Limnohabitans_A | LimnohabitanA sp021300495 |
| envZ: two-component system, OmpR family, osmolarity sensor histidine kinase EnvZ | K07638 | 4 | yellow | Pseudomonadota | Gammaproteobacteria | Burkholderiales | Methylophilaceae | BACL14 | BACL14 sp018607785 |
| envZ: two-component system, OmpR family, osmolarity sensor histidine kinase EnvZ | K07638 | 0 | grey | Pseudomonadota | Gammaproteobacteria | Burkholderiales | Rhodocyclaceae | Sulfuritalea | |
| osmB: osmotically inducible lipoprotein OsmB | K04062 | 0 | grey | Pseudomonadota | Gammaproteobacteria | Burkholderiales | Rhodocyclaceae | Sulfuritalea | |
| envZ: two-component system, OmpR family, osmolarity sensor histidine kinase EnvZ | K07638 | 0 | grey | Pseudomonadota | Gammaproteobacteria | Burkholderiales | Rhodocyclaceae | Zoogloea | Zoogloea sp016720685 |
| opuBD: osmoprotectant transport system permease protein | K05846 | 0 | grey | Pseudomonadota | Gammaproteobacteria | Burkholderiales | SG8-41 | Ga0077527 | |
| osmB: osmotically inducible lipoprotein OsmB | K04062 | 0 | grey | Pseudomonadota | Gammaproteobacteria | Burkholderiales | SG8-41 | Ga0077527 | |
| opuBD: osmoprotectant transport system permease protein | K05846 | 0 | grey | Pseudomonadota | Gammaproteobacteria | Pseudomonadales | Halomonadaceae | Halomonas | Halomonas campaniensis |
| opuA: osmoprotectant transport system ATP-binding protein | K05847 | 0 | grey | Pseudomonadota | Gammaproteobacteria | Pseudomonadales | Halomonadaceae | Halomonas | Halomonas campaniensis |
| opuC: osmoprotectant transport system substrate-binding protein | K05845 | 0 | grey | Pseudomonadota | Gammaproteobacteria | Pseudomonadales | Halomonadaceae | Halomonas | Halomonas campaniensis |
| opuA: osmoprotectant transport system ATP-binding protein | K05847 | 0 | grey | Pseudomonadota | Gammaproteobacteria | Pseudomonadales | Pseudohongiellaceae | UBA9145 | UBA9145 sp003483155 |
| opuBD: osmoprotectant transport system permease protein | K05846 | 0 | grey | Pseudomonadota | Gammaproteobacteria | Steroidobacterales | Steroidobacteraceae | RPQJ01 | |
| opuA: osmoprotectant transport system ATP-binding protein | K05847 | 0 | grey | Pseudomonadota | Gammaproteobacteria | Steroidobacterales | Steroidobacteraceae | RPQJ01 | |
