## Supplemental Figures for "Suspended and sinking particle-associated microbiomes exhibit distinct lifestyles in the Elbe estuary"

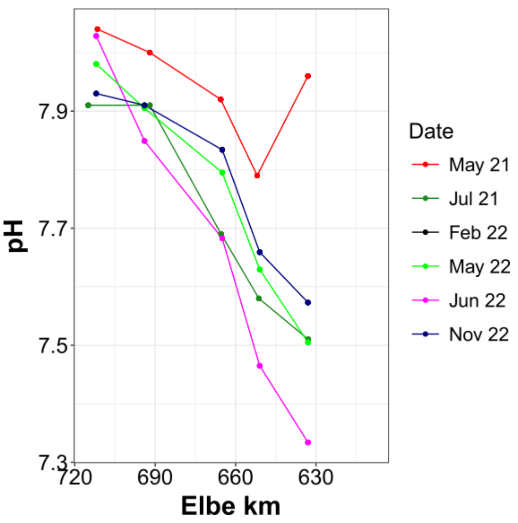


**Figure S1. pH changes over the Elbe Estuary.** Colours denote sample dates from May 21 to Nov 22.

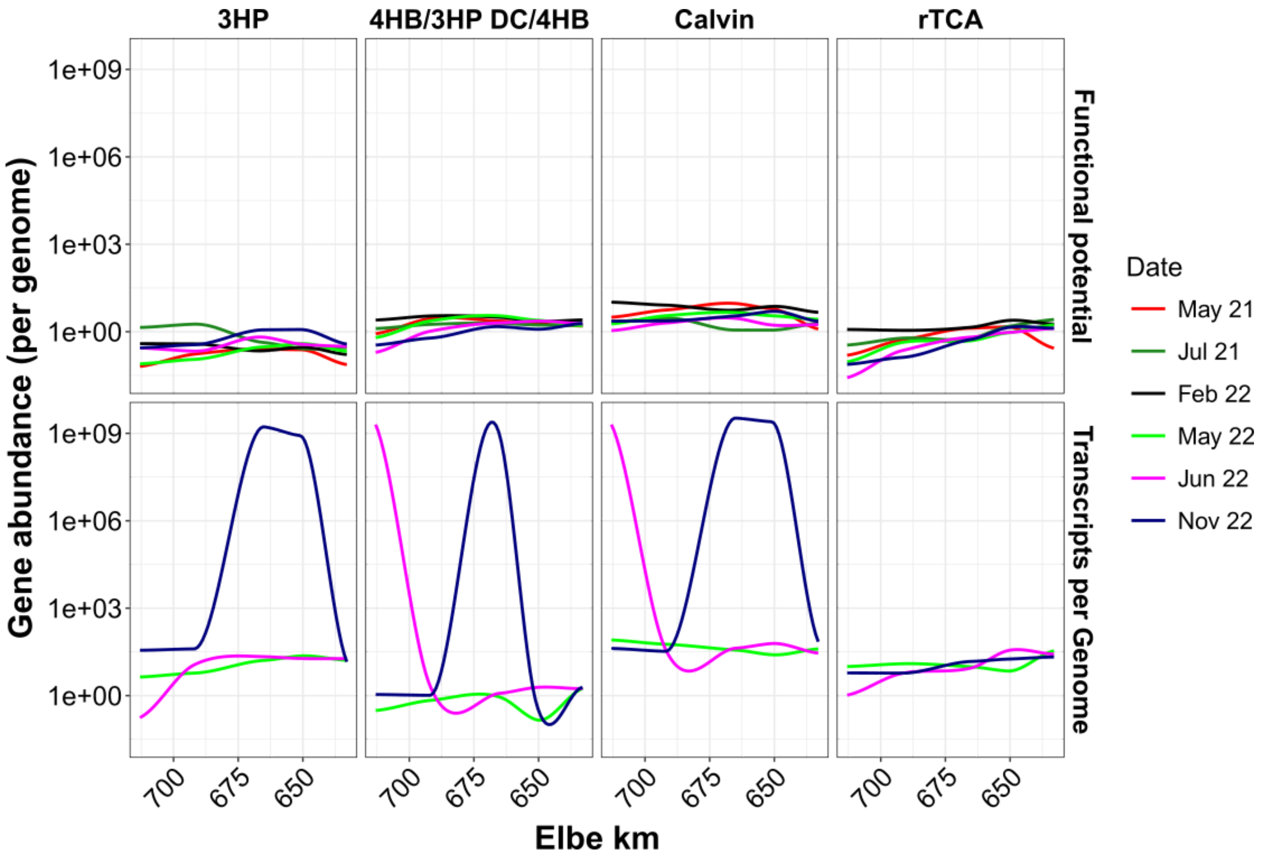


**Figure S2. Carbon fixation pathway functional potential and transcription in the Elbe Estuary.** The mean of two samples is depicted using different colours for sample dates. Carbon fixation pathways are the reverse tricarboxylic acid (rTCA) cycle, the 3-hydroxypropionate (3HP) bi-cycle, the

4-hydroxybutyrate/3-hydroxypropionate (4HB/3HP) cycle, the dicarboxylate/4-hydroxybutyrate (DC/4HB) cycle, the reductive acetyl-CoA pathway (Wood–Ljungdahl pathway—WLP), and the reductive glycine pathway.

Since WLP and reductive glycine pathways operate in both oxidative and reductive directions with identical enzymes (e.g., in methane or acetate oxidation), we are unable to predict the direction and cannot proceed with further analysis.

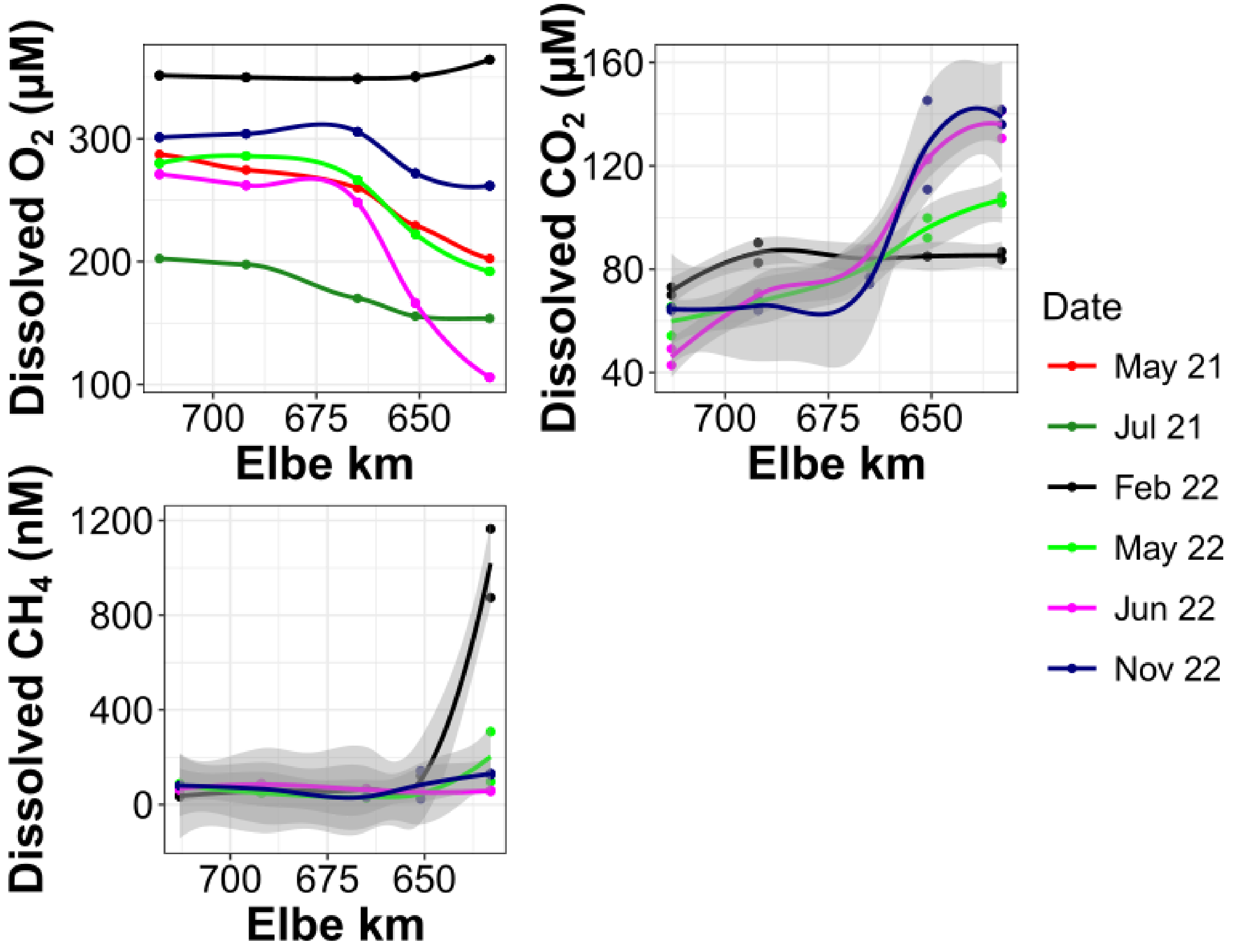


**Figure S3. Dissolved gases in the Elbe Estuary.** Dissolved O2, CO2, and CH4 are shown across the Elbe Estuary, with colour noting sampling dates. Shaded areas represent the 95% confidence interval.

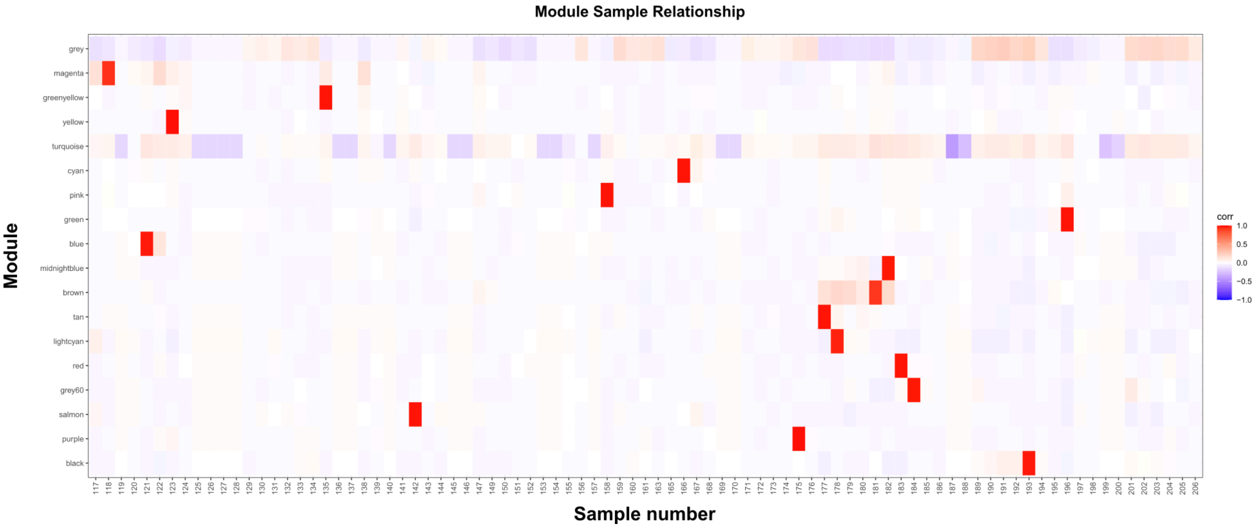


**Figure S4. Transcripts per Gene network modules and the influence of individual samples.** Colour denotes the Pearson correlation strength.

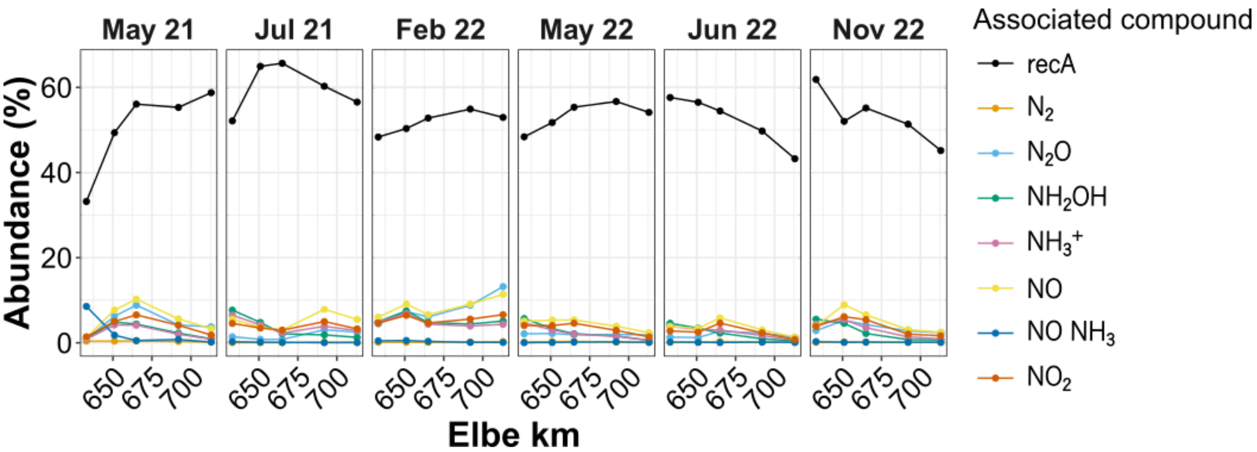


**Figure S5. Nitrogen cycling associated gene abundance.** The mean of two samples abundance (cell normalised) of genes associated with the processing of the colour dependent compounds.

**Table S1. Osmotic pressure associated genes identified in WGCNA module related MAGs.**
